# Global analysis of protein degradation in prion infected cells

**DOI:** 10.1101/712927

**Authors:** Charles Hutti, Kevin A. Welle, Jennifer R. Hryhorenko, Sina Ghaemmaghami

## Abstract

Prion diseases are rare neurological disorders caused by the misfolding of the cellular prion protein (PrP^C^) into cytotoxic fibrils (PrP^Sc^). Intracellular PrP^Sc^ aggregates primarily accumulate within late endosomes and lysosomes, organelles that participate in the degradation and turnover of a large subset of the proteome. Thus, intracellular accumulation of PrP^Sc^ aggregates have the potential to globally influence protein degradation kinetics within an infected cell. We analyzed the proteome-wide effect of prion infection on protein degradation rates in N2a neuroblastoma cells by dynamic stable isotopic labeling with amino acids in cell culture (dSILAC) and bottom-up proteomics. The analysis quantified the degradation rates of more than 4,700 proteins in prion-infected and uninfected cells. As expected, the degradation rate of the prion protein is significantly decreased upon aggregation in infected cells. In contrast, the degradation kinetics of the remainder of the N2a proteome generally increases upon prion infection. This effect occurs concurrently with increases in the cellular activities of autophagy and lysosomal hydrolases. The resulting enhancement in proteome flux may play a role in the survival of N2a cells upon prion infection.

## INTRODUCTION

Prion diseases are infectious neurodegenerative disorders caused by the cytotoxic aggregation of the prion protein (PrP)^1-3^. These diseases are present in a number of mammalian species and include scrapie in sheep, bovine spongiform encephalopathy in cattle and chronic wasting disease in deer. In humans, prion diseases include Kuru, Creutzfeldt-Jakob Disease, Fatal Familial Insomnia, and Gerstmann-Straussler-Scheinker Syndrome^4,5^. During the course of each of these diseases, aggregates of the prion protein (PrP^Sc^) accumulate and cause the conversion of the non-toxic cellular form of the prion protein (PrP^C^) into additional PrP^Sc^ in a self-seeding fashion^6^. While the function of PrP^C^ remains controversial, PrP^Sc^ is thought to be the sole cause of prion diseases.

In uninfected cells, the prion protein resides primarily on the cell surface and is degraded through the endocytic pathway as part of its natural turnover cycle^7,8,9^. While PrP^C^ is readily degraded by lysosomal hydrolases, PrP^Sc^ appears to be more resistant to proteolysis and accumulates within endosomal and lysosomal compartments during the course of infection^7,10,11^. Over time, PrP^Sc^ accumulates intracellularly and eventually results in cell death and dispersion of aggregates into the extracellular space^12^. Prion aggregates can cause the conversion of PrP^C^ on adjacent cells, thus spreading PrP^Sc^ aggregates throughout the brain. While the exact mechanism of pathogenicity is unknown, the accumulation of PrP^Sc^ in the lysosome and endosomal vesicles is thought to contribute to the cytotoxicity of prions^13^.

The lysosome serves as a proteolytic center for two major protein degradation pathways within the cell: autophagy and endosome mediated degradation^14,15^. Lysosomal vesicles facilitate the hydrolysis of proteins into their constitutive amino acids by subjecting them to low pH unfolding conditions and acidic hydrolases. Given that PrP^Sc^ accumulates in the lysosome during the course of prion infection, several studies have investigated the direct and indirect effects of accumulating prion aggregates on lysosomal degradation^13^. In mouse brains infected with prion aggregates, the expression of lysosomal hydrolases are upregulated at the level of transcription^16^. Similarly, in cultured N2a cells, the activities of lysosomal hydrolases have been shown to be upregulated during prion infection^17^. In both mouse and hamster brains infected with prion aggregates, there are increases in the abundances of autophagy-related proteins p62 and LC3–II^18^. While increases in the abundances of p62 and LC3-II can also be interpreted as markers of late-stage autophagic inhibition, N2a cells exhibit increased expression of p62 mRNA, supporting the hypothesis that autophagy is upregulated upon prion infection. Together, these studies suggest that host cells upregulate lysosomal degradation as a response to prion infection. However, other studies have provided evidence suggesting that accumulation of prion aggregates inhibit lysosomal degradation by interfering with the ability of the cell to endocytose and degrade proteins^19^. Considering these opposing effects of prion infection on intracellular protein degradation, the net effect of PrP^Sc^ accumulation on global protein turnover kinetics remains unclear. Thus, we sought to utilize a proteomic approach to conduct a global analysis of changes in protein degradation kinetics during prion infection.

Recent advances in quantitative mass spectrometry and bottom-up proteomics have enabled the measurement of protein turnover kinetics on a global scale^20-22^. These studies have shown that protein half-lives can range from minutes to years and can be influenced by several intrinsic and extrinsic factors. As examples, the presence of specific degradation sequence motifs (degrons), as well as a protein’s physical properties such as isoelectric points, surface area, thermodynamic stabilities, and molecular weights can influence a protein’s inherent half-life^23-28^. Additionally, the relative activities of specific protein degradation pathways such as autophagy and the ubiquitin proteasome system (UPS) can influence the degradation kinetics of specific subsets of the proteome^29-32^. By using modern proteomic techniques such as dynamic stable isotopic labeling of amino acids in cell culture (dSILAC), the *in vivo* degradation rate of individual proteins within the proteome can be measured in different cellular and environmental conditions^32,33^. In this study, we used dSILAC to investigate the proteome-wide impact of intracellular PrP^Sc^ accumulation on protein clearance kinetics in prion-infected cells.

## RESULTS

### Measurement of degradation kinetics by dSILAC

An overview of the dSILAC methodology used in this study is illustrated in Figure 1A and details of the experimental design are described in Materials and Methods. Briefly, cells are cultured in media containing isotopically labeled amino acids (^13^C-lysines and ^13^C-arginines). The rate by which unlabeled (“light”) proteins within the cell are replaced by newly synthesized labeled (“heavy”) proteins can be quantified by mass spectrometry over time. The quantified rate constant for fractional labeling is commonly referred to as the protein clearance rate (*k*_*clearance*_)^21,34^. The measurement of *k*_*clearance*_ can be conducted on proteome-wide scales using tandem mass spectrometry and a bottom-up proteomic workflow^35^. In dividing cells, measured rates of protein clearance represent the additive effects of two factors: protein degradation (*k*_*degradation*_) and protein dilution due to cell division (*k*_*division*_) (Figure 1B). Thus, degradation rate constants can be obtained by subtracting experimentally determined rates of cell division from rates of clearance^21,36^.

**Figure 1.**
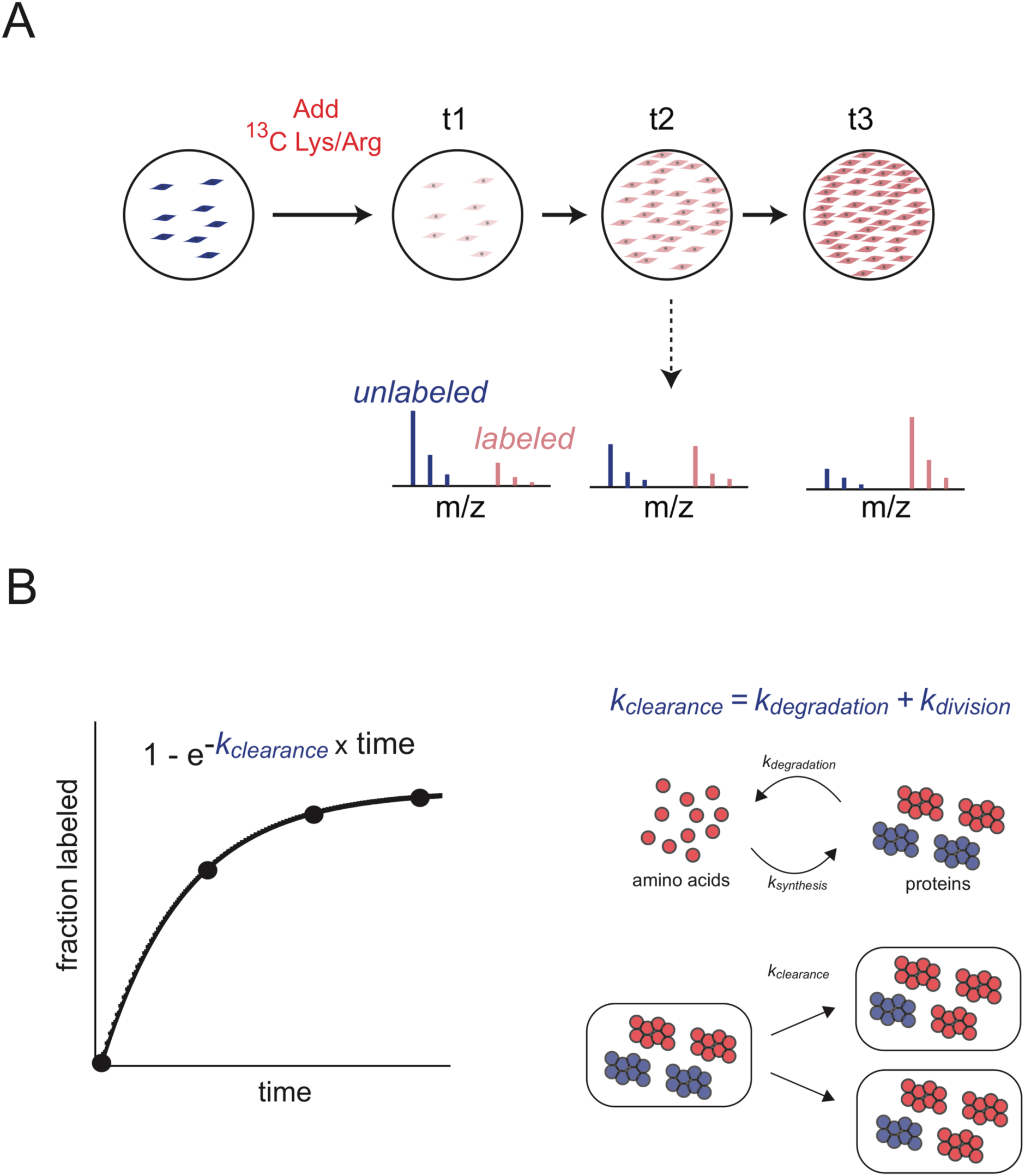
Experimental design and quantitative analysis of dynamic stable isotopic labeling of amino acids in cell culture (dSILAC) experiments. A) Experimental design. Cultured cells are grown in the presence of ^13^C-lysine and ^13^C-arginine. Newly synthesized proteins incorporate these heavy labeled amino acids as the original pool of unlabeled proteins is cleared over time. The kinetics of fractional labeling of proteins is measured by LC-MS/MS analysis using a bottom-up proteomics workflow. B) Quantitative analysis. The first order rate constant of fractional labeling is a measure of the protein clearance rate. Two factors contribute to the rate of clearance: the rate of dilution due to cell division and the rate of protein degradation. The blue and red colors respectively represent unlabeled and labeled amino acids, proteins, spectra and cells.

### Prion infected and uninfected cultured cell models

For our analysis, we utilized a transgenic clone of N2a neuroblastoma cells that highly overexpress the PrP gene (N2a-Cl3) ^37,38^. When infected with the Rocky Mountain Laboratories (RML) prion strain, these cells accumulate protease resistant PrP^Sc^ at cellular levels that are equivalent to terminal prion infected brain tissues^37^. As a transformed cell line, N2a cells are known to have an unstable karyotype. Thus, to obtain an isogenic uninfected control, prion infected cells were treated with the antiprion compound quinacrine^39,40^. We cleared populations of infected N2a-Cl3 cells of PrP^Sc^ by treating them with a standard dosage of quinacrine for four weeks and confirmed the clearance of protease-resistant PrP^Sc^ by western blots (Figure 2A). Prion-cleared controls were propagated in the absence of quinacrine for an additional week prior to isotopic labeling in order to remove any potentially confounding effects of quinacrine in the media. Prion infected and cleared N2a-Cl3 cells are referred to as −QA and +QA in this study.

**Figure 2.**
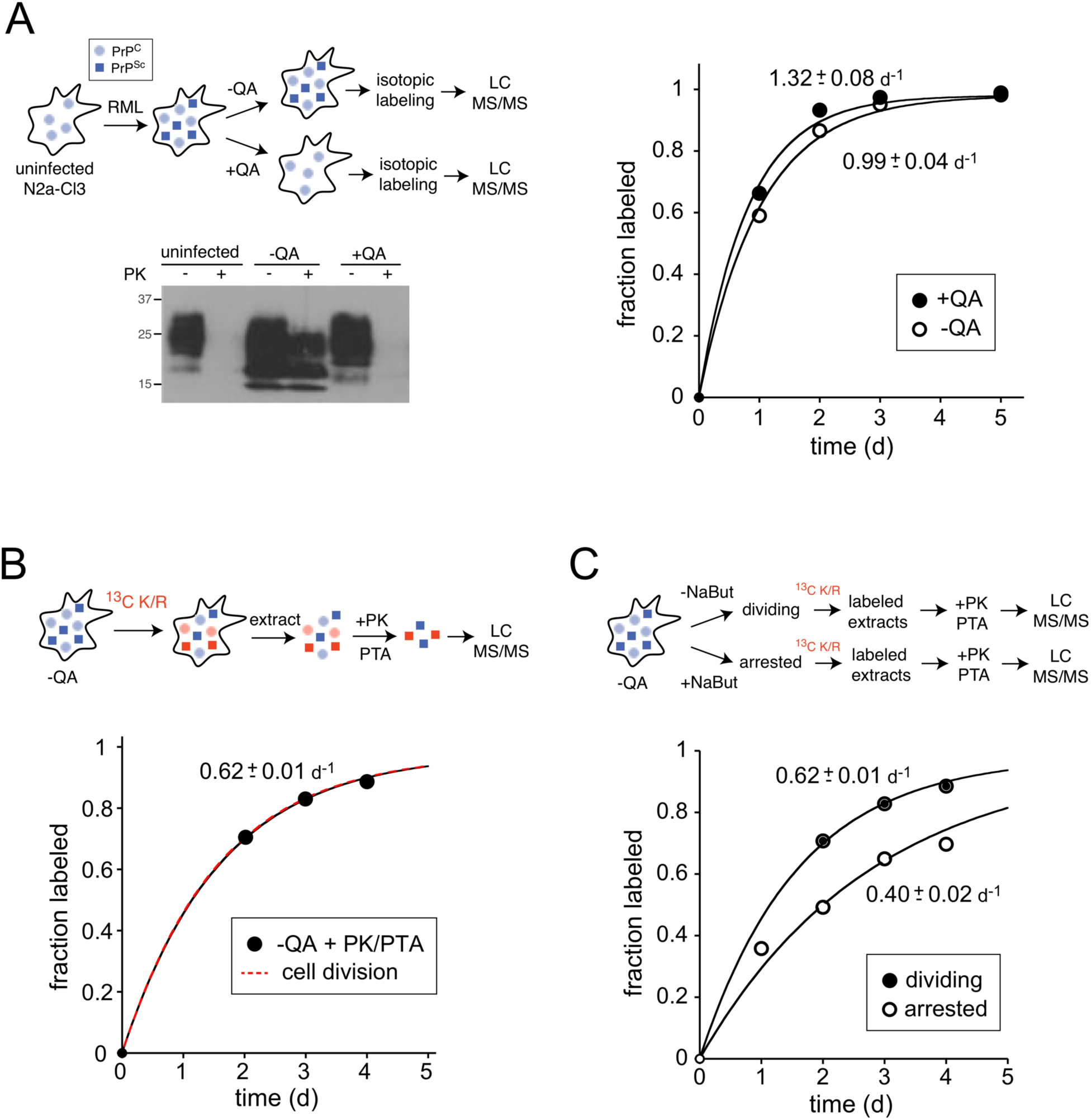
The clearance kinetics of PrP^C^ and PrP^Sc^. A) Clearance kinetics of total PrP. N2a-Cl3 cells were infected with the RML prion strain (−QA). Some passaged cultures were treated with quinacrine to generate an uninfected control (+QA). The presence and absence of protease-resistant PrP^Sc^ in −QA and +QA samples were verified by western blots following PK digestion. −QA and +QA cultures were propagated in labeling media containing ^13^C Lysine/Arginine, harvested at different time points and analyzed by LC-MS/MS as described in Figure 1. The kinetic plots indicate the average fractional labeling of all peptides mapped to PrP (PrP^C^ for +QA and PrP^C^ plus PrP^Sc^ for −QA). B) Clearance kinetics of PrP^Sc^. Lysates from −QA cells were subjected to proteinase K digestion to isolate protease resistant PrP^Sc^ prior to LC-MS/MS analysis. Clearance kinetics were analyzed as in (A). C) The effect of cell division on the clearance rate of PrP^Sc^. To determine the contribution of cell division to PrP^Sc^ clearance, a population of - QA cells was treated with sodium butyrate to halt cell division prior to dSILAC analysis. Clearance kinetics were analyzed as in (A). The results indicate that PrP^Sc^ aggregates are cleared more rapidly in dividing cells.

### The clearance kinetics of the prion protein in −QA and +QA cells

We conducted dSILAC analyses on −QA and +QA cells and were able to quantify the clearance kinetics of 58,590 peptides mapped to 5,215 proteins in −QA cells and 59,320 peptides mapped to 5,168 proteins in +QA cells (Table 1). 4,730 proteins were shared between the two datasets. We initially focused on analyzing the kinetic data mapped to PrP. We observed that *k*_*clearance*_ of the total prion protein population was faster in +QA cells in comparison to −QA cells (Figure 2A). Measured *k*_*clearance*_ values for PrP in +QA and −QA cells were 1.32 d^−1^ and 0.99 d^−1^, respectively. The degradation rate of PrP^C^ in dividing cells was calculated as 0.70 d^−1^ by subtracting the rate of cell division from the clearance rate. The slower rate of PrP clearance in −QA cells is consistent with the fact that PrP^Sc^ aggregates are partially resistant to cellular proteolysis and thus have a slower degradation rate than PrP^C^.

**Table 1.** Coverage of dSILAC experiments

| Cells | Peptides identified | Proteins identified | Peptides quantified | Proteins quantified | Median $k_{deg}$ (d<sup>-1</sup>) | Shared proteins |
| --- | --- | --- | --- | --- | --- | --- |
| -QA | 64,648 | 7,280 | 58,590 | 5,215 | 0.27 | 4,730 |
| +QA | 63,355 | 7,196 | 59,320 | 5,168 | 0.24 |  |

However, this analysis is complicated by the fact that both PrP^C^ and PrP^Sc^ are present in −QA cells, and peptides from both populations are simultaneously quantified and contribute to the observed fractional labeling. Thus, in order to measure the degradation rate of PrP^Sc^ alone, a second dSILAC experiment was performed where lysates were treated with proteinase K and the protease-resistant PrP^Sc^ was isolated by phosphotungsic acid (PTA) precipitation^41^ prior to LC-MS/MS analysis (Figure 2B). PrP data from this experiment established the *k*_*clearance*_ of PrP^Sc^ in dividing N2a cells as 0.62 d^−1^. Importantly, this rate of clearance exactly mirrors the measured rate of cell division in these cells (Figure 2B). This observation suggests that the degradation of PrP^Sc^ in dividing prion infected cells is inhibited to such an extent that its clearance occurs almost entirely by cellular dilution rather than degradation.

If prion clearance in dividing cells occurs primarily by dilution due to cell division, then the arrest of cell division should substantially decrease the observed *k*_*clearance*_ of PrP^Sc^. To test this hypothesis, a third dSILAC experiment was performed where cell division was arrested 48 hours prior to the introduction of ^13^C lysine/arginine by the addition of sodium butyrate, which has been shown to arrest cell division and induce the differentiation of N2A cells to neuron-like cells^42-45^. Labeled extracts from sodium butyrate-treated division-arrested cells were treated with proteinase K and protease-resistant PrP^Sc^ was isolated by PTA precipitation prior to LC-MS/MS analysis (Figure 2C). As predicted, we observed that the clearance rate of PrP^Sc^ in division arrested cells (0.40 d^−1^) is significantly slower than that found in dividing cells. Together, this data confirms that the clearance of PrP in prion infected cells is substantially slowed upon formation of PrP^Sc^ aggregates and validates dSILAC as a methodology capable of quantitatively analyzing changes protein clearance kinetics in N2a-Cl3 cells.

### Global effect of PrP^Sc^ accumulation on proteome turnover

The data obtained from the dSILAC experiment described in Figure 2A were used to analyze the proteome-wide effect of PrP^Sc^ aggregates on protein clearance (Supplementary Tables S1-3). We limited our analysis to 4,730 proteins where heavy to light ratios (H/L) could be quantified for two or more peptides in at least three timepoints in both −QA and +QA samples (Table 1). In order to determine the true degradation rates (*k*_*degradation*_) of proteins, the rate of cell division was subtracted from the measured rate of clearance (*k*_*clearance*_) Figure 3 provides a comparison of protein *k*_*degradation*_ values between −QA and +QA cells as histograms, pairwise scatter plot and log_2_ ratios. Globally, it is evident that unlike PrP itself, prion infection does not result in dramatic reductions in degradation rates of most proteins in N2a-Cl3 cells. Indeed, we observed a slight but statistically significant proteome-wide increase in degradation rates in −QA cells (Figure 3B, 3C).

**Figure 3.**
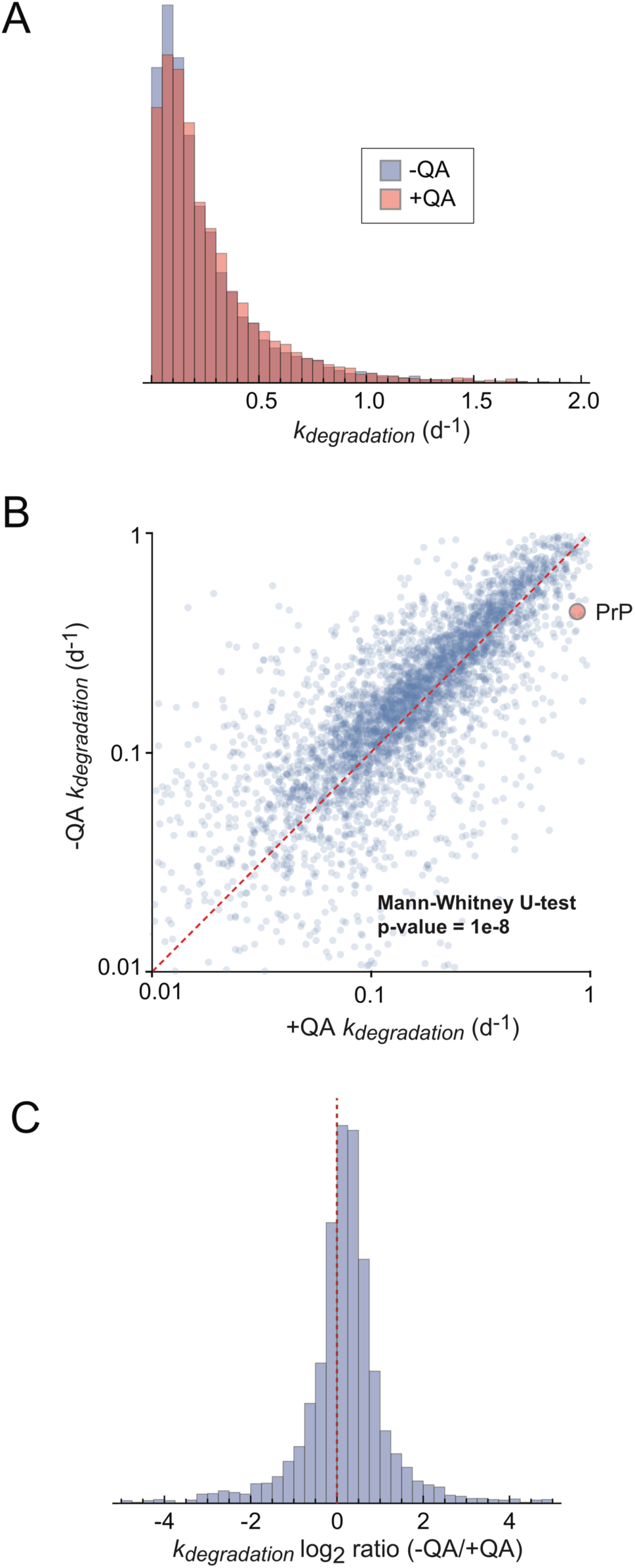
Global analysis of protein degradation rates. Using dSILAC, degradation rates were measured for 4,730 proteins shared between −QA and +QA samples. The rates are compared as distribution plots (A), pairwise comparisons (B) and log2 ratios (C). In (B), the dotted line indicates the identity line and red circle indicates the comparison of PrP^C^ and PrP^Sc^ in −QA and −QA samples, respectively. The −QA and +QA datasets differ with a p-value of 1e-8 using the Mann Whitney U-test.

The proteome-wide data supports the hypothesis that the accumulation of prion aggregates results in the upregulation of cellular degradation machinery. Interestingly, we observed that the increase in degradation rates is disproportionately evident in long-lived proteins, defined as proteins with *k*_*degradation*_ values less than 0.2 d^−1^ (Figure 4A). In eukaryotic cells, the two main protein degradation pathways with general selectivity are the UPS and lysosomal pathways, such as autophagy. It is generally accepted that short-lived proteins are degraded by the UPS, whereas long-lived proteins are degraded by the lysosome^31,46^. We therefore explored the possibility that prion infection results in the upregulation of the autophagy pathway, resulting in an increase in the degradation rate of autophagic substrates.

**Figure 4.**
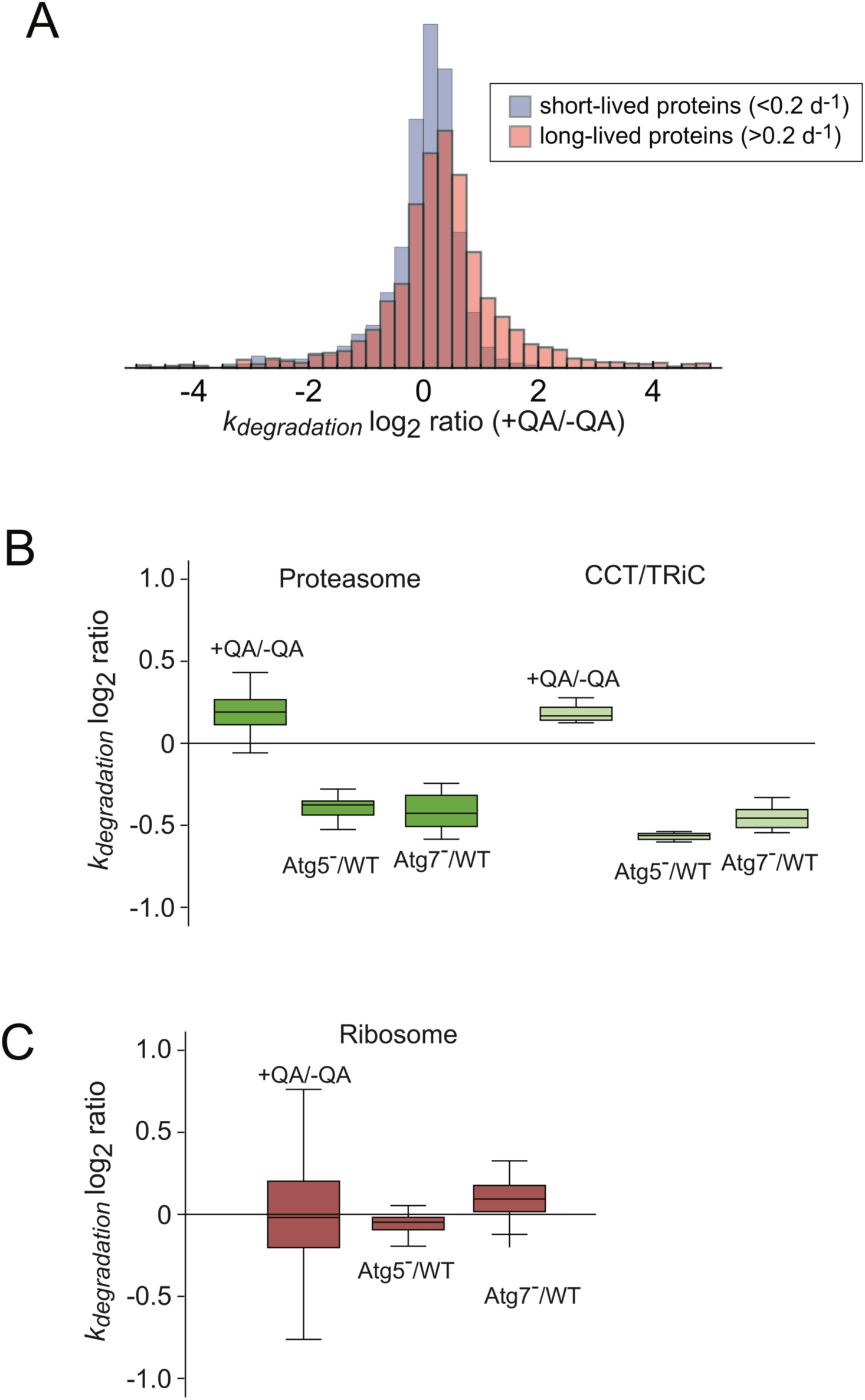
The degradation rates of long-lived proteins and autophagy substrates are increased in prion infected cells. A) The log_2_ ratio of degradation rates between −QA and +QA samples for short-lived (k_degradation_ > 0.2 d^−1^) and long-lived (k_degradation_ < 0.2 d^−1^) proteins. B) The effect of prion infection on k_degradation_ of two previously established substrates of basal autophagy: the proteasome and CCT/TRiC. The change in degradation rates in autophagy-deficient cells (ATG5^−/−^ and ATG7^−/−^) compared to wildtype, measured by Zhang et al., is shown for comparison. C) The effect of prion infection on k_degradation_ of the ribosome, previously shown to be excluded from basal autophagy by Zhang et al.

In a previous study, we conducted a proteome-wide dSILAC study to identify proteins whose degradation rates are diminished in cells lacking two core components of canonical autophagy: Atg5 and Atg7^32^. The study identified the subunits of the 26S proteasome and the CCT/TRiC chaperonin as *bona fide* substrates of basal autophagy. As a counter-example, the degradation rate of the ribosome was unaltered in Atg5 and Atg7 knockout backgrounds, indicating that it was excluded from basal autophagy. We determined how the degradation rates of these complexes are impacted by prion infection (Figure 4B-C). Our data indicate that the degradation of the proteasome and CCT/TRiC are increased in −QA cells in comparison to +QA cells. Conversely, the degradation rate of ribosomal subunits remained unchanged. This observation is consistent with the idea that the rate of autophagy is enhanced in prion infected cells.

### Enhancement of autophagy in prion infected cells

Using western blots, we analyzed −QA and +QA extracts for a number of reporters associated with autophagy and lysosomal degradation (Figure 5A). These included Cathepsins D, L and A (lysosomal hydrolases), LC3-II (marker of autophagsomes) and p62/SQSTM1 (autophagy substrate and reporter of autophagic flux)^47,48^. We observed an increase in the level of LC3-II relative to LC3-I, indicating an increase in the steady-state levels of autophagosomes in prion infected cells. Consistent with the upregulation of autophagic flux, our dSILAC results indicate that p62/SQSTM1 degradation rates are increased in prion infected cells (Figure 5B). Interestingly, despite its enhanced degradation, the steady-state levels of p62/SQSTM1 are not drastically diminished in prion infected cells (Figure 5A). This observation is consistent with previous reports showing that N2a cells exhibit increased expression of p62 mRNA upon prion infection^18^. Additionally, we observed that levels of some, but not all, lysosomal cathepsins are greatly enhanced in prion infected cells (Figure 5A). This effect was particularly dramatic for cathepsin D. To verify this finding, total cathepsin D enzymatic activity was assayed in +QA and −QA using a synthetic fluorescent substrate. The data indicated that the total activity of cathepsin D is significantly increased in prion infected cells (Figure 5C). To determine whether this increase in activity may be due to overall expansion of lysosomal compartments, the total lysosomal volume within +QA and −QA cells were quantified using a fluorometric lysosomal stain (CytoPainter). The data indicated that there is a significant expansion of total lysosomal volume in prion infected cells (Figure 5D). Together, the data suggest that prion infection is associated with the upregulation of specific molecular components of lysosomal degradation. Specifically, prion infection in N2a cells is associated with expansion of lysosomal compartments and enhanced delivery of protein targets to lysosomes resulting from the activation of the autophagy pathway.

**Figure 5.**
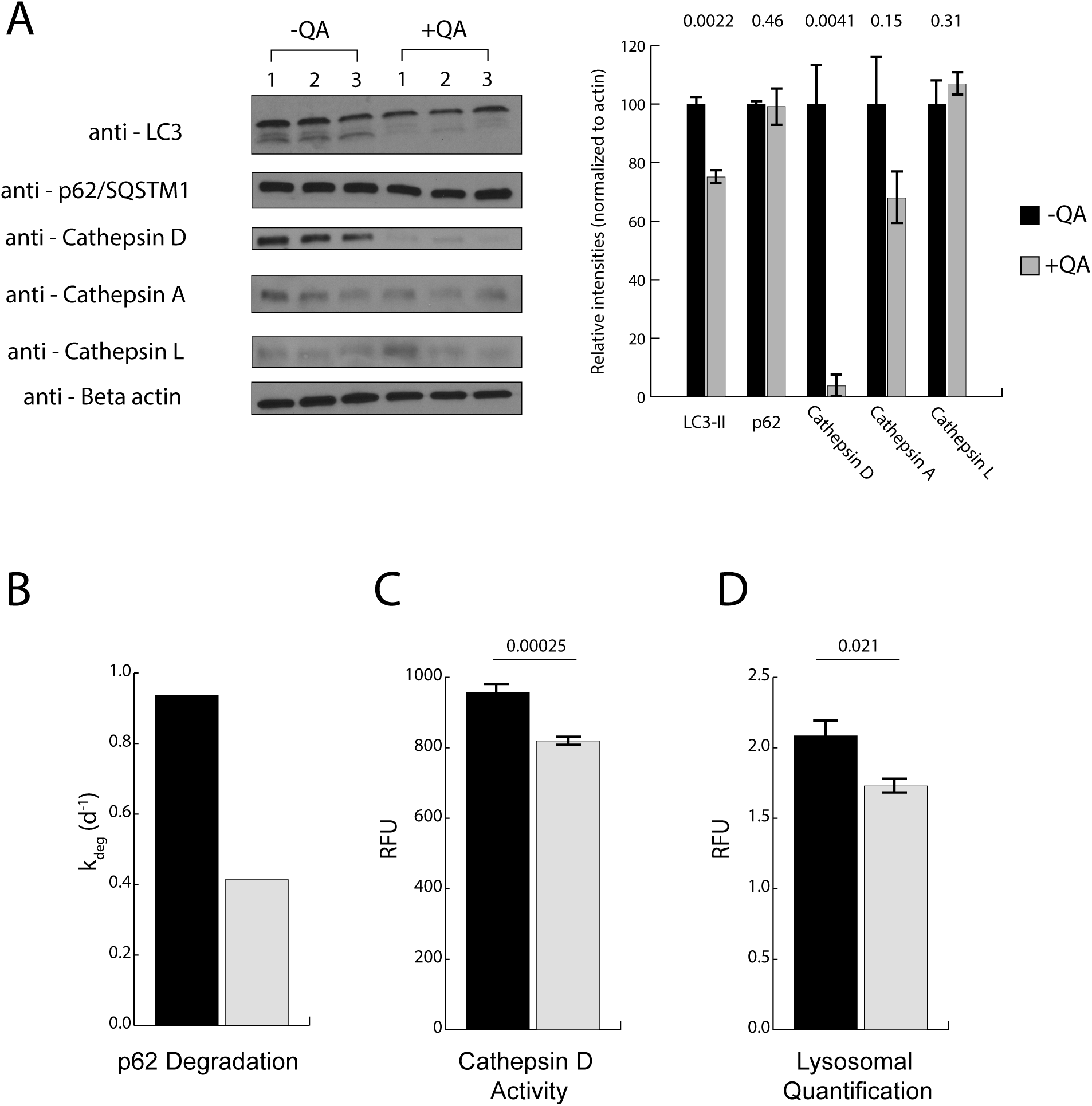
Biochemical analysis of markers of lysosomal degradation. A) Western blot analysis of −QA and +QA protein lysates (done in biological triplicate) indicate significant increases in LC3-II and cathepsin D abundance in −QA cells, whereas little change is observed in levels of Cathepsins A, L and p62/SQSTM1. Similarly, the abundance of the major lysosomal hydrolase cathepsin D is significantly increased in −QA cells, consistent with increases to protein degradation. B) While the abundance p62 is not significantly altered during prion infection (A), the degradation rate of p62 (determined by dSILAC analysis) is significantly increased. C) Consistent with increases in its steady-state levels (A), the total enzymatic activity of Cathepsin D is increased in prion infected −QA cells. D) Total lysosomal volume, determined by a fluorometric lysosomal stain, is increased in prion infected −QA cells. The error bars indicate standard error of the mean. Comparisons between −QA and +QA samples were conducted by two-tailed Student t-test. Numbers above the bar graphs indicate the calculated p-value.

## DISCUSSION

The cytopathology of prion diseases is known to be intimately linked to the lysosomal degradation system. Expansion of lysosomes, late endosomes and autophagic vesicles are well-established neuropathological features of prion infected cells both *in vivo* and *in vitro*^10,49,50^. Delivery of PrP^Sc^ to lysosomes by autophagy is a key pathway for its degradation, and levels of intracellular PrP^Sc^ is readily modulated by inhibition or activation of autophagy^51,52^. During prion infection, the conversion of PrP^C^ to PrP^Sc^ results in its intracellular stabilization where it accumulates in late endosomes and lysosomes^7,11^. The accumulation of PrP^Sc^ aggregates in lysosomal compartments has the potential to influence lysosomal degradative pathways that are normally responsible for turnover of a significant fraction of the cellular proteome. Accordingly, a number of studies have sought to characterize the impact of prion infection on protein degradation and cellular proteostasis. In a recent study, Homma et al. showed that prion infection impairs the degradative capacity of lysosomes by interfering with their maturation^18^. Other studies have shown that autophagy is upregulated in prion infected tissues and cells, suggestive of a possible compensatory response to restore proteostasis upon accumulation of PrP^Sc 10,49,50^. However, in these studies, changes in protein degradation were monitored for a small number of markers and individual proteins, making it difficult to assess the proteome-wide effects of prion infection on protein degradation. To overcome this limitation, we used a dSILAC strategy to quantify changes in proteome degradative flux in prion infected cells. Our results provided a number of insights regarding the effects of prion infection on the degradation kinetics of PrP and the proteome at large.

Consistent with previous studies, our results indicate that the degradation rate of PrP^Sc^ in N2a-Cl3 cells is significantly slower than PrP^C^. Our measured degradation rates for PrP^C^ and PrP^Sc^ were 0.70 d^−1^ (half life = 1.0 d) and 0.40 d^−1^ (half life = 1.7 d), respectively. The measured degradation rate for PrP^Sc^ is consistent with rates previously measured in a bigenic mouse model (half-life = 1.5 d)^53^ and ScN2a cells measured by radiolabeled pulse-chase experiments (half-life > 1d)^8^. The measured degradation rate for PrP^C^, although comparable to that reported for bigenic mice (half life = 18 h)^53^, is significantly slower than that previously reported for N2a cells (half life = 5 h)^8^. A number of factors may account for this discrepancy. First, in this study, we have used a transgenic overexpression variant of N2a cells. The increased expression level of PrP may influence its clearance kinetics by formation of non-infectious aggregates, as has been previously reported for some mouse overexpression lines^54^ or through partial inhibition of protein degradation pathways. Alternatively, this discrepancy may be due to differences in the methodology used for measurement of degradation rates. In previously conducted pulse-chase analyses, newly synthesized proteins were labeled with short pulses of ^35^ S-methionine and their degradation kinetics were measured following an unlabeled chase^8^. Here, our methodology involved continuous labeling experiments where the turnover of the entire protein pool (not just newly synthesized proteins) was monitored over time. It has recently been shown that the turnover of many proteins is multiphasic, and a large fraction of newly synthesized proteins are rapidly degraded before becoming incorporated into the steady-state protein pool^55^. Thus, short-term pulse-chase experiments may underestimate the half-life of the steady-state protein pool. Regardless, the relative stabilization of PrP^Sc^ that was reported in previous studies was recapitulated in our experiments.

Importantly, our results indicate that, in prion infected cells, PrP^Sc^ is sufficiently stabilized such that its dominant route of clearance in dividing ScN2a cells is cellular dilution through cytokinesis rather than proteolytic degradation. These results are consistent with previous results showing that PrP^Sc^ levels increase in cultured cells when they reach a state of confluency and there is a reduction in the division rate^56^. In this way, the clearance of PrP^Sc^ in dividing cultured cells fundamentally differs from postmitotic neurons *in vivo*, where the catabolism of PrP^Sc^ is the sole route of clearance.

Our data indicate that the impact of prion infection on global protein degradation is generally subtle. The median prion-induced change in the degradation rate of the proteome is ∼10% (Figure 3, Table 1). Nonetheless, there is a statistically significant enhancement of degradation rates for long-lived proteins, including at least two protein complexes (the proteasome and CCT/TRiC chaperonin) that are established substrates of basal autophagy^32^. Conversely, the degradation of short-lived proteins known to be enriched in substrates of UPS^31^, and the ribosome which has been shown to be excluded from basal autophagy^32^ are minimally impacted by prion infection. Consistent with previous studies^16,18^, we further showed that levels of LC3-II, a marker of autophagosomes, and expression levels of specific lysosomal hydrolases are increased in prion infected cells. This upregulation was accompanied by an overall expansion of lysosomal compartments in the cell. Together, the data provide evidence that that the rate of autophagic flux is enhanced in N2a cells in response to prion infection.

Together, our results highlight two potential mechanisms that may enable dividing N2a cells to maintain viability in a prion infected state. First, the process of cell division acts as a continuous clearance mechanism, reducing the steady-state level of PrP^Sc^ in dividing cells. Second, the activation of autophagy may partially mitigate the proteostatic disruptions caused by the formation and accumulation of PrP^Sc^. By enhancing the rate of autophagic flux, the cells stimulate a degradation system that may counter the stability of PrP^Sc^ and enhance the turnover of other autophagic substrates. Thus, as has been shown in a number of *in vivo* and *in vitro* experiments^13^, further enhancement of autophagy through pharmacological induction of lysosomal activity may provide a promising strategy for clearance of PrP^Sc^ and maintaining proteostasis in prion-infected cells.

## MATERIALS AND METHODS

### Cell culture

Cell cultures of the mouse neuroblastoma, N2a-Cl3, were maintained in Eagle’s Minimum Essential Medium (ATCC) supplemented with 15% FBS (Invitrogen), 100 U/mL penicillin, 100 U/mL streptomycin at 37 °C with 5% CO_2_.

### Generation of isogenic controls

N2a-Cl3 cells previously infected with the Rocky Mountain Laboratories strain of prion aggregate (RML) were treated with 4uM of quinacrine and cleared of prion infection within four passages.

### Stable isotope labeling

The media utilized for isotopic labeling was Eagle’s minimum essential medium (ATCC) supplemented with 15% dialyzed FBS (Thermo Scientific), 100 U/mL penicillin, and 100 U/mL streptomycin. Cells were gradually adapted to this media by replacing normal FBS with dialyzed FBS within five passages. Cells were then plated at a density of 1,000,000 cells per 10-cm plate.

One day after plating, the dividing cultures were switched to MEM labeling media for SILAC (Thermo Scientific) supplemented with L-arginine:HCl (^13^C6, 99%) and L-lysine:2HCl (^13^C6, 99%; Cambridge Isotope Laboratories) at concentrations of 0.1264 g/L and 0.087 g/L and 15% dialyzed FBS (Thermo Scientific), 100 U/mL penicillin, and 100 U/mL streptomycin. For whole proteome analysis, cells were collected after 0, 1, 2, 3, and 5 d of labeling and washed with PBS. For analysis of isolated PrP^Sc^ aggregates, cells were collected after 0, 1, 2, 3, and 4 d of labeling and washed with PBS. All cell pellets were frozen before further analysis.

### Mass spectrometry sample preparation

Cells were lysed with ice-cold lysis buffer (10 mM Tris·HCl pH 8.0, 0.15 M NaCl, 0.5% Nonidet P-40, 0.48% SDS). Cell lysates were centrifuged at 16,000 × g for 10 min and the supernatants were then transferred to new Eppendorf tubes. Protein concentrations were measured by the bicinchoninic assay (BCA) kit (Thermo Scientific).

When processing lysates for prion protein peptides, 20 µg of proteinase K (fungal) (Thermo Scientific) was added to 1 mg cell lysates in 1 mL of lysis buffer. Lysates were incubated for 1 hour at 37 °C. To halt further ribosome translation, 20 µL of PMSF was added to lysates for a final concentration of 20 mM. Sodium lauroyl sarcosinate was added for a final 1% weight to volume concentration, and 75 µL of Phosophotungsic acid (pH 7.4) was added for a final 0.75% weight to volume to precipitate aggregated proteins from the lysate. Lysates were incubated for 3 hours at 37 °C while shaking at 350 rpm. Lysates were then centrifuged at 16,000 × g for 30 minutes and the supernatants were then transferred to new Eppendorf tubes, leaving 30 µL of lysate left to resuspend pelleted aggregates. 12.5 µL of resuspended pellet was added to 12.5 µL of 5% SDS and boiled for 5 minutes before processing further processing.

Protein lysates after proteinase K digestion as well as 25 µg of total protein from whole cell extracts were processed into peptides by the following S-trap protocol. Reduction of disulfide bonds was performed with 5 mM Tris(2-carboxyethyl)phosphine (TCEP) Bond-breaker (Thermo Scientific) at RT for 1 h, and protein alkylation was performed with 10 mM iodoacetamide (IAA) at RT for 30 min in darkness. DTT was added to 1 mM to quench IAA and samples were applied to S-Trap Micro Spin Columns (Protifi). To derive tryptic peptides, 20 µL of 50ng/ µL trypsin (selective cleavage on the C-terminal side of lysine and arginine residues) was added and the samples were incubated overnight at 37 °C in a water bath. Peptides were released from the column using subsequent washes of 0.1% TFA followed by 50% ACN in 0.1% TFA.

To increase proteome coverage, high-pH fractionation was conducted on whole cell extracts before LC-MS/MS using homemade C18 spin columns. Samples were dried down and resuspended in 50 µL of 100 mM ammonium formate (pH 10). Eight different elution buffers were made in 100 mM ammonium formate (pH 10) with 5%, 7.5%, 10%, 12.5%, 15%, 17.5%, 20%, and 50% acetonitrile added. After conditioning the column with acetonitrile and 100 mM ammonium formate, the samples were added and centrifuged. An ammonium formate wash was performed to remove any residual salt before the eight elutions were collected in fresh tubes. All samples were then dried down and re-suspended in 15 µL of 0.1% TFA.

### LC-MS/MS analysis

#### Q Exactive Plus LC-MS/MS

Peptides were injected onto a homemade 30 cm C18 column with 1.8 um beads (Sepax), with an Easy nLC-1000 HPLC (Thermo Fisher), connected to a Q Exactive Plus mass spectrometer (Thermo Fisher). Solvent A was 0.1% formic acid in water, while solvent B was 0.1% formic acid in acetonitrile. Ions were introduced to the mass spectrometer using a Nanospray Flex source operating at 2 kV. Optimized gradients for individual fractions were employed, as shown in Supplementary Table S4. The Q Exactive Plus was operated in data-dependent mode, with a full scan followed by 20 MS/MS scans. The full scan was done over a range of 400-1400 m/z, with a resolution of 70,000 at m/z of 200, an AGC target of 1e6, and a maximum injection time of 50 ms. Peptides with a charge state between 2-5 were picked for fragmentation. Precursor ions were fragmented by higher-energy collisional dissociation (HCD) using a collision energy of 27 and an isolation width of 1.5 m/z, with an offset of 0.3 m/z. MS2 scans were collected with a resolution of 17,500, a maximum injection time of 55 ms, and an AGC setting of 5e4. Dynamic exclusion was set to 25 seconds.

#### Fusion Lumos LC-MS/MS

The HPLC and ion source set-up was identical to the Q Exactive, except that the nLC was a 1200 series with 80% acetonitrile in 0.1% formic acid as solvent B. The gradient began at 3% B and held for 2 minutes, increased to 10% B over 5 minutes, increased to 38% B over 38 minutes, then ramped up to 90% B in 3 minutes and was held for 3 minutes, before returning to starting conditions in 2 minutes and re-equilibrating for 7 minutes, for a total run time of 60 minutes. The Fusion Lumos was operated in data-dependent mode, while also employing an inclusion list containing SILAC labeled PrP peptides that was created using the Skyline software program (Supplementary Table S5). An MS2 scan even would be triggered when a precursor ion was within 10 ppm of a m/z value on the inclusion list. Otherwise, the method proceeded as usual. The cycle time was set to 1.5 seconds. Monoisotopic Precursor Selection (MIPS) was set to Peptide. The full scan was done over a range of 375-1400 m/z, with a resolution of 120,000 at m/z of 200, an AGC target of 4e5, and a maximum injection time of 50 ms. Peptides with a charge state between 2-5 were picked for fragmentation. Precursor ions from the inclusion list and those selected by the data-dependent method were fragmented by collision induced dissociation (CID) using a collision energy of 30% and an isolation width of 1.1 m/z. MS2 scans were collected with the ion trap scan rate set to rapid, a maximum injection time of 200 ms, and an AGC setting of 1e4. Dynamic exclusion was set to 20 seconds.

### Data analysis and kinetic models

MS2 data for all samples were searched against the *M. musculus* (22,305 entries, downloaded 8/7/2017) uniprot databases using the integrated Andromeda search engine with MaxQuant software. Peptide searches were done as previously described ^36^ with the addition of the match-between-runs with a match time window of 0.7 minutes, a match ion mobility window of 0.05, an alignment time window of 20 minutes, and an alignment ion mobility of 1 (Supplementary Table S6).

The determination of degradation rate constants (*k*_*degradation*_) from fraction labeled measurements were conducted in accordance to the kinetic model outlined previously.^36^

### Western blotting

Cells were lysed with ice-cold lysis buffer (10 mM Tris·HCl pH 8.0, 0.15 M NaCl, 0.5% Nonidet P-40, 0.48% SDS). Cell lysates were centrifuged at 16,000 × g for 10 min and the supernatants were then transferred to new Eppendorf tubes. For Western blot analysis, 20 μg was separated by electrophoresis in 10% polyacrylamide gels and transferred to polyvinylidene difluoride membranes using Trans-Blot Turbo Transfer System (Biorad). After 1 h incubation at room temperature in Odyssey Blocking Buffer in TBS (Licor), the membrane was incubated with the indicated antibodies at 4 °C overnight. The membranes were then washed with TBST/0.1% Tween 20, and the corresponding secondary antibodies were applied to the membranes for 1 h at room temperature. The membranes were then washed with TBST/0.1% Tween 20, and the detection of signal was done using an enhanced chemiluminescence detection kit (Pierce). The primary antibodies and the corresponding dilutions utilized for the Western blots were Anti beta-Actin Antibody: 1:5000 (Abcam); Anti-p62 (SQSTM1) Antibody: 1:1,000 (MBL International); Anti-LC3 Antibody: 1:1,000 (MBL International); Cathepsin A (A-19): 1:1000 (Santa Cruz Technology), Cathepsin D (c-20): 1:1000 (Santa Cruz Technology), Cathepsin L: 1:1000 (Santa Cruz Technology), and anti-PrP (D18) 1:1000 (compliments of the Pruisner lab). In instances where multiple proteins of varying molecular weights were analyzed from the same SDS/PAGE gel, the membrane was cut into multiple parts following transfer. Each part was probed with a different primary antibody against a specific target protein and, subsequently, with the corresponding secondary antibody.

### Lysosome activity analysis

Cathepsin D activity (used as a proxy for total lysosomal hydrolase activity) was measured by using Cathepsin D Activity Assay Kit (Abcam). Cells were collected, and 350 μL of lysis buffer was used to lyse 1 million cells; 5 μL of lysate was incubated in substrate/buffer solution for 75 minutes at 37 °C. Enzyme activity was measured by monitoring release of the fluorescent cleavage product, MCA. Activity measurements from parallel reactions containing 0.7 μM protease inhibitor pepstatin A were subtracted from the activity measurements obtained without the inhibitor. Fluorescence was quantified by SpectraMax M5 at Ex/Em = 328/460 nm.

### Lysosomal Quantification

Lysosomal quantity was measured using the Lysosomal Staining Kit – Green Fluorescence – Cytopainter (Abcam). Cells were plated at 60,000 cells per well of a 96 well plate. One day after plating, media was replaced with the dye working solution and incubated for 60 minutes at 37 °C. Parallel reactions containing 10 nM of the lysosomal inhibitor Bafilomycin were subtracted from the activity measurements obtained without the inhibitor. Fluorescence was quantified by SpectraMax M5 at Ex/Em = 490/525 nm.

## Data availability

All raw and processed data are available at ProteomeXchange Consortium via the PRIDE database (www.edi.ac.uk/pride/; accession number PXD014577).

## Supplementary Figures

Table S1 – Peptide level search data

Table S2 – Protein level search data

Table S3 – Protein degradation rates

Table S4 – Optimized gradients for individual fractions during QExactive LC-MS/MS

Table S5 – Inclusion list for SILAC labeled PrP peptides generated using Skyline software program

Table S6 – MaxQuant software parameters for data analysis

## Notes

https://www.edi.ac.uk/pride/

