## Supplementary figures and images for "Global analysis of protein degradation in prion infected cells"

### Supplemental Table 4

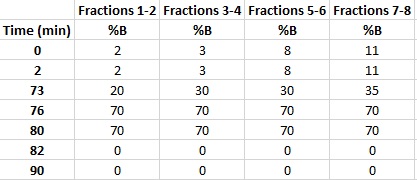
